# Structure and computation-guided design of a mutation-integrated trimeric RBD candidate vaccine with broad neutralization against SARS-CoV-2

**DOI:** 10.1101/2021.06.18.448958

**Authors:** Yu Liang, Jing Zhang, Run Yu Yuan, Mei Yu Wang, Peng He, Ji Guo Su, Zi Bo Han, Yu Qin Jin, Jun Wei Hou, Hao Zhang, Xue Feng Zhang, Shuai Shao, Ya Nan Hou, Zhao Ming Liu, Li Fang Du, Fu Jie Shen, Wei Min Zhou, Fang Tang, Ze Hua Lei, Shuo Liu, Wei Zhen, Jin Juan Wu, Xiang Zheng, Ning Liu, Shi Chen, Zhi Jing Ma, Fan Zheng, Si Yu Ren, Zhong Yu Hu, Gui Zhen Wu, Wei Jin Huang, Chang Wen Ke, Qi Ming Li

## Abstract

The spike (S) protein receptor-binding domain (RBD) of SARS-CoV-2 is an attractive target for COVID-19 vaccine developments, which naturally exists in a trimeric form. Here, guided by structural and computational analyses, we present a mutation-integrated trimeric form of RBD (mutI tri-RBD) as a broadly protective vaccine candidate, in which three RBDs were individually grafted from three different circulating SARS-CoV-2 strains including the prototype, Beta (B.1.351) and Kappa (B.1.617). The three RBDs were then connected end-to-end and co-assembled to possibly mimic the native trimeric arrangements in the natural S protein trimer. The recombinant expression of the mutI tri-RBD, as well as the homo-tri-RBD where the three RBDs were all truncated from the prototype strain, by mammalian cell exhibited correct folding, strong bio-activities, and high stability. The immunization of both the mutI tri-RBD and homo-tri-RBD plus aluminum adjuvant induced high levels of specific IgG and neutralizing antibodies against the SARS-CoV-2 prototype strain in mice. Notably, regarding to the “immune-escape” Beta (B.1.351) variant, mutI tri-RBD elicited significantly higher neutralizing antibody titers than homo-tri-RBD. Furthermore, due to harboring the immune-resistant mutations as well as the evolutionarily convergent hotspots, the designed mutI tri-RBD also induced strong broadly neutralizing activities against various SARS-CoV-2 variants, especially the variants partially resistant to homo-tri-RBD. Homo-tri-RBD has been approved by the China National Medical Products Administration to enter clinical trial (No. NCT04869592), and the superior broad neutralization performances against SARS-CoV-2 support the mutI tri-RBD as a more promising vaccine candidate for further clinical developments.

## Introduction

Severe acute respiratory syndrome coronavirus 2 (SARS-CoV-2), which causes the respiratory disease of COVID-19, has spread rapidly around the world since the end of 2019 and resulted in a pandemic of this new coronavirus infections^1,2^. SARS-CoV-2 is highly pathogenic and virulent in human, which presents a great threat to public health and imposes a heavy burden on global economy. To date, significant progresses have been achieved in COVID-19 vaccine developments, and several vaccines have been authorized by World Health Organization (WHO) for emergency use^3–5^, which is encouraging to eliminate the threat of COVID-19. However, SARS-CoV-2 evolves continuously and many new variants have emerged, among which four strains, including the Alpha (B.1.1.7), Beta (B.1.351), Gamma (P.1) and Delta (B.1.617.2), have been classified as variants of concern (VOC) by the World Health Organization (WHO). Besides that, other six variants have been categorized as variants of interest (VOI) according to the weekly reports of the WHO on Jun 8, 2021^6^. More and more evidences have demonstrated that many of the variants, especially those of VOC, exhibit increased transmissibility and virulence. More seriously, the mutations in some of these variants may also potentially lead to immune escape, which may reduce the effectiveness of existing vaccines and therapeutics^7^. The B.1.351 and P.1 variants have been proved by multiple evidences to be able to evade from the neutralizing antibodies elicited by natural infections and vaccinations^8^. The ChAdOx1 nCoV-19 vaccine has been proved in clinical trials to be unprotective against the mild-to-moderate COVID-19 caused by the B.1.351 variant^9^. Several preliminary studies also suggested that the B.1.617 variant is less sensitive to the neutralization by the sera from the vaccinated individuals and resistant to some monoclonal antibodies^10^. Exhaustive analysis of the available SARS-CoV-2 sequences has revealed that several immune-resistant mutations in the circulating variants were also evolutionarily convergent hotspots^11^, indicating the possible recurrence of these mutations individually or combinedly in the future. The possible immune escape of the currently-circulating as well as future-emerging SARS-CoV-2 variants has raised concerns about the efficacy of existing vaccines, and also raised the urgent requirements to develop vaccines with broad protections against various virus strains^12^. A commonly used strategy for the development of multi-protective vaccine is to produce multivalent vaccine through the simple combination of monovalent vaccines against individual virus stains. But this strategy is time-consuming and cost-expensive. Another more promising, also more difficult, method to develop multivalent vaccine is the construction of hybrid vaccine via integrating multiple circulating variants into one vaccine. Here, adopting the second strategy, we present a mutation-integrated trimeric form of receptor binding domain (mutI tri-RBD) as a broadly protective vaccine candidate, which combined the antigens derived from different variants into one vaccine and simultaneously integrated various key mutations into a single immunogen.

In SARS-CoV-2, the spike (S) glycoproteins decorated on the surface of the virion are responsible for recognition of the host cell receptor, i.e. human angiotensin converting enzyme 2 (hACE2), to mediate virus entry into the cell. The S protein is naturally self-assembled into a homotrimer anchored in the viral membrane, and each monomer is composed of two subunits, S1 and S2^13,14^. The receptor binding domain (RBD) in S1 subunit is protruding from the viral surface and directly involved in the interactions with the receptor hACE2, which serves as a key target for COVID-19 vaccine development. In addition, many immunological investigations also showed that S1 RBD contains multiple neutralizing epitopes, which is immunodominant in eliciting neutralizing antibody responses to SARS-CoV-2 infection^15–17^. It has been revealed that S1 RBD-based vaccine candidates can elicit neutralizing antibody responses against both the pseudo and live SARS-CoV-2 in vitro, and also induce protective immunity in non-human primates in vivo^18,19^. However, due to the small molecular size of RBD monomer as the immunogen, monomeric RBD-based vaccines are challenged by their limited immunogenicity^20,21^. A potential strategy to improve the immunogenicity is polymerization of RBDs to increase the molecular size and generate antigen with multiple copies of antigenic determinant. Dai et al. recently produced a dimeric form of SARS-CoV-2 S1 RBD through tandemly connecting two RBDs into a single molecule, which exhibits significantly improved immunogenicity with neutralizing antibody titers 10- to 100-fold higher than those induced by the conventional S1 RBD monomer^20,21^. However, in the native structure S protein is self-assembled into a trimer and thus S1 RBDs also naturally exist in a homo-trimeric form. It is reasonably speculated that the construction of trimeric form of RBD resembling its native conformational arrangements could achieve better immunogenic properties. To our knowledge, no trimerized RBD-based COVID-19 subunit vaccine is reported until now. Structural analysis of the S trimer indicates that the N- and C- termini of the RBD are close to each other and there exists a long loop with high flexibility at both termini, which potentially allow the formation of RBD trimer (tri-RBD) through end-to-end connection to possibly mimic the RBD arrangements in the natural S protein trimer. Furthermore, the tri-RBD that accommodates three RBDs in one molecule enables us to construct a multivalent vaccine candidate, in which three RBDs are individually derived from different circulating SARS-CoV-2 variants and co-assembled into a hybrid trimer. Therefore, guided by structural and computational analyses, a mutI tri-RBD was designed as a vaccine candidate with broadly protective capability, in which three RBDs from different SARS-CoV-2 strains were connected end-to-end and co-assembled to possibly mimic the native trimeric arrangements in the natural S protein trimer. As a comparison, a homo-tri-RBD, in which three RBDs were all derived from the prototype SARS-CoV-2 strain, was also constructed. Then, the biochemical characterizations and immunological properties of the recombinant mutI tri-RBD were evaluated comprehensively and compared with those of homo-tri-RBD, which support the mutI tri-RBD as a promising vaccine candidate for further clinical developments.

## Results and Discussion

### Structure and computation-guided construction of the mutI tri-RBD as an immunogen

In the native state, S protein is assembled into homo-trimer, and the three RBDs in the S trimer are also naturally arranged in a trimeric form. We speculate that the trimeric form of RBD resembling its naturally adopted state should exhibit better immunogenicity than the monomeric or dimeric RBDs, due to the larger molecular size and multiple copies of the antigenic determinant. In addition, to elicit broadly protective immunity against the various emerging SARS-CoV-2 strains, the development of multivalent vaccine through possible incorporation of multiple immunodominant RBDs from different circulating variants into a single immunogenic molecule is a promising, though difficult, strategy. At present, neither trimeric form of RBD-based COVID-19 vaccine nor broadly protective vaccine against various SARS-CoV-2 strains has been reported to our knowledge. Therefore, guided by the analysis of the native three-dimensional structure of S protein, we sought to construct a possible hybrid trimeric form of RBD, i.e., mutI tri-RBD, as a COVID-19 vaccine candidate with broad protections against various virus strains.

We present a mutI tri-RBD through the end-to-end connection of three RBDs derived from different circulating SARS-CoV-2 variants to possibly mimic the RBD arrangements in the natural S protein trimer. This RBD trimer design strategy was proposed mainly based on the following considerations: (1) In S protein, the N- and C- termini of the RBD are close to each other, which enables the multimerization of the RBDs through end-to-end connections without serious steric clashes. (2) There exist long flexible loops at both termini of the RBD. Although the termini of different RBDs are rather far apart in the native structure of S trimer, the flexible terminal loops may facilitate the connection of these RBDs without destroying the core structure of individual RBD. (3) The tri-RBD construction strategy enables the realization of multivalent vaccine via a hybrid connection of three different RBDs, which are individually derived from different circulating strains of SARS-CoV-2, into a single immunogenic molecule. From the native tertiary structure of S protein, a RBD truncation scheme that meets the above considerations were designed, as shown in Extended Data Fig. 1a. The truncation scheme comprises the residues 319-537, which contains the entire RBD with long loops at both the N- and C- termini. Besides that, according to the epidemic trends and the capability of immune escape, the RBDs from the prototype, Beta (B.1.351) and Kappa (B.1.617.1) strains, respectively, were selected for the hybrid construction of multivalent immunogen. Compared with the prototype strain, the RBD of B.1.351 variant harbors three residue mutations, i.e., K417N, E484K and N501Y, and the RBD of B.1.617.1 contains L452R and E484Q mutations, respectively. Several of these mutations were also revealed to be evolutionarily convergent. The three truncated RBDs were then connected end-to-end to generate mutI tri-RBD, and a homo-tri-RBD constructed only based on the early prototype strain was also produced as a comparison (Figure 1a).

**Fig. 1.**
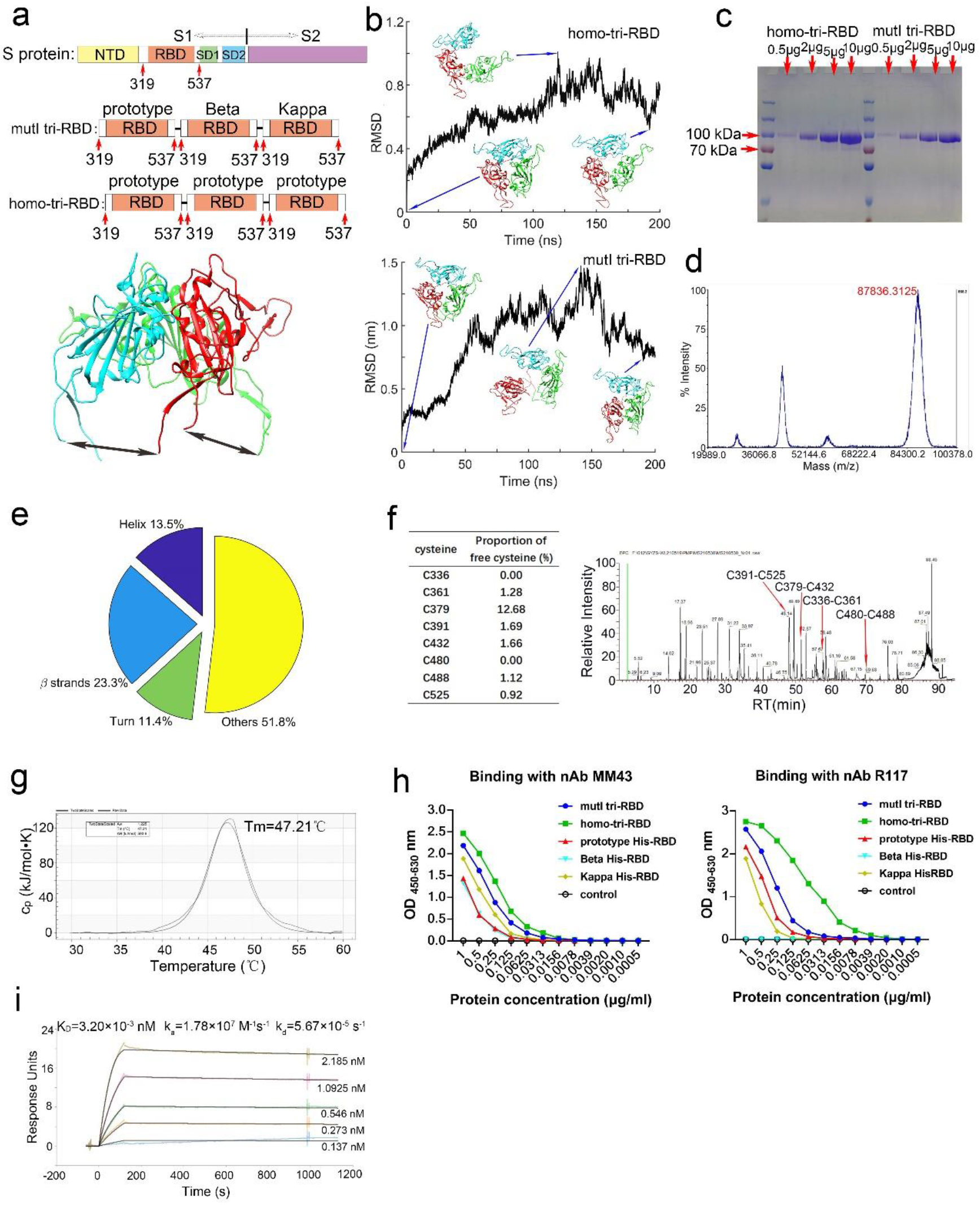
Structure-guided design, production, and characterization of the mutI tri-RBD. **a.**A schematic illustration of the mutI tri-RBD and homo-tri-RBD design schemes. The RBD region comprising the residues 319-537 was truncated from the S protein, and three truncated RBDs were connected end-by-end to construct the trimeric forms of RBD. In mutI tri-RBD, three RBDs were individually derived from three different circulating SARS-CoV-2 strains, i.e., the prototype, Beta (B.1.351) and Kappa (B.1.617.1). In homo-tri-RBD, the three RBD units were all truncated from the prototype strain. In the upper subfigure, the S1 and S2 subunits of the S protein, as well as NTD, RBD, SD1 and SD2 in S1 subunit, are marked. The lower subfigure displays the natural trimeric arrangement of RBDs in the native structure of S trimer. The arrows indicate the direct connections of the N- and C- terminals between different RBDs. **b.**Structural modeling and MD simulation of the designed homo-tri-RBD (upper subfigure) and mutI tri-RBD (lower subfigure). Time-evolution of the C_α_ root-mean square deviation (RMSD) of the modelled structure during MD simulation, as well as several snapshot conformations in the simulation, is displayed. **c.**SDS-PAGE profiles of increasing amounts of the recombinant mutI tri-RBD and homo-tri-RBD proteins expressed by HEK293T cells. **d.**Molecular weight of mutI tri-RBD determined by MALDI-TOF MS. **e.**Secondary structure contents of mutI tri-RBD protein analyzed by circular dichroism spectrometry. **f.**Left: The proportions of free sulfhydryl for all the cysteine residues in mutI tri-RBD. Right: The disulfide-linkages in the recombinant mutI tri-RBD protein detected by liquid chromatography-mass spectrometry. Only the disulfide bonds in one RBD unit are listed. **g.**Differential scanning calorimetry thermograms of the recombinant mutI tri-RBD protein. **h.**The binding capability of the designed mutI tri-RBD and homo-tri-RBD proteins with two anti-RBD monoclonal nAbs, i.e., MM43 and R117, evaluated by ELISA. As controls, the binding activities with the monoclonal nAbs for the monomeric his-tagged RBDs from the prototype, Beta (B.1.351) and Kappa (B.1.617.1) SARS-CoV-2 strains were also measured. **i.**The binding profiles of the recombinant mutI tri-RBD with hACE2 detected by surface plasmon resonance assay.

Using the native structure of S trimer as the template, the possible tertiary structures of the designed mutI tri-RBD and homo-tri-RBDs were built by Modeller9.23 software^22^. To analyze the stability of the modelled structures as well as the intrinsic conformational motions encoded in the structures, the constructed trimeric structures were also subjected to 200 ns all-atomic MD simulations with GROMACS software^23^. The structural modelling and MD simulation results showed that the modelled mutI tri-RBD and homo-tri-RBD structures were stereo-chemically reasonable and rather stable without serious steric clashes, and one RBD in the modelled trimeric structure exhibits an intrinsic “down” to “up” hinge-like motion during MD simulations that is similar to the conformational transition of the RBD in the natural S protein (Figure 1b as well as Supplementary Movie 1 and 2). It should be mentioned that we do not exclude the possibility that the connected three RBDs assembled into other trimeric form different from that in the native structure of S trimer, which needs further validation by the experimental determination of its three-dimensional structure in the future research.

### Expression, purification, and characterization of the recombinant mutI tri-RBD and homo-tri-RBD

The designed mutI tri-RBD as well as homo-tri-RBD proteins of SARS-CoV-2 were transiently expressed by using the mammalian HEK293T cells. The culture supernatant of the transfected cells was harvested and purified by chromatography combined with ultrafiltration. SDS-PAGE exhibited an obvious single band with the molecular mass about 90 kDa both for the recombinant mutI tri-RBD and homo-tri-RBD proteins (Figure 1c), which indicates the well-expression of the trimeric form of RBD for the designed two schemes. The purity of the prepared proteins is over 95% both for mutI tri-RBD and homo-tri-RBD. Furthermore, for mutI tri-RBD protein, the exact molecular weight was determined to be around 87836 Da by matrix assisted laser desorption ionization-time of flight mass spectrometer (MALDI-TOF MS) (Figure 1d), which is larger than the theoretical value of about 74 kD calculated based on the amino acid sequence. The difference between the experimental and calculated molecular weights implies the dense glycosylation of the protein. To characterize the glycosylation of the recombinant mutI tri-RBD protein, the N- and O-glycosylation sites were detected and analyzed by using liquid chromatography-mass spectrometry (LC-MS) and the BioPharma Finder data analyzing software. Multiple possible N-glycosylation sites on asparagine and O-glycosylation sites on serine and threonine were detected (Extended Data Fig. 2 and 3), and the locations of these identified glycosylation sites were distributed widely over the surface of RBD.

**Fig. 2.**
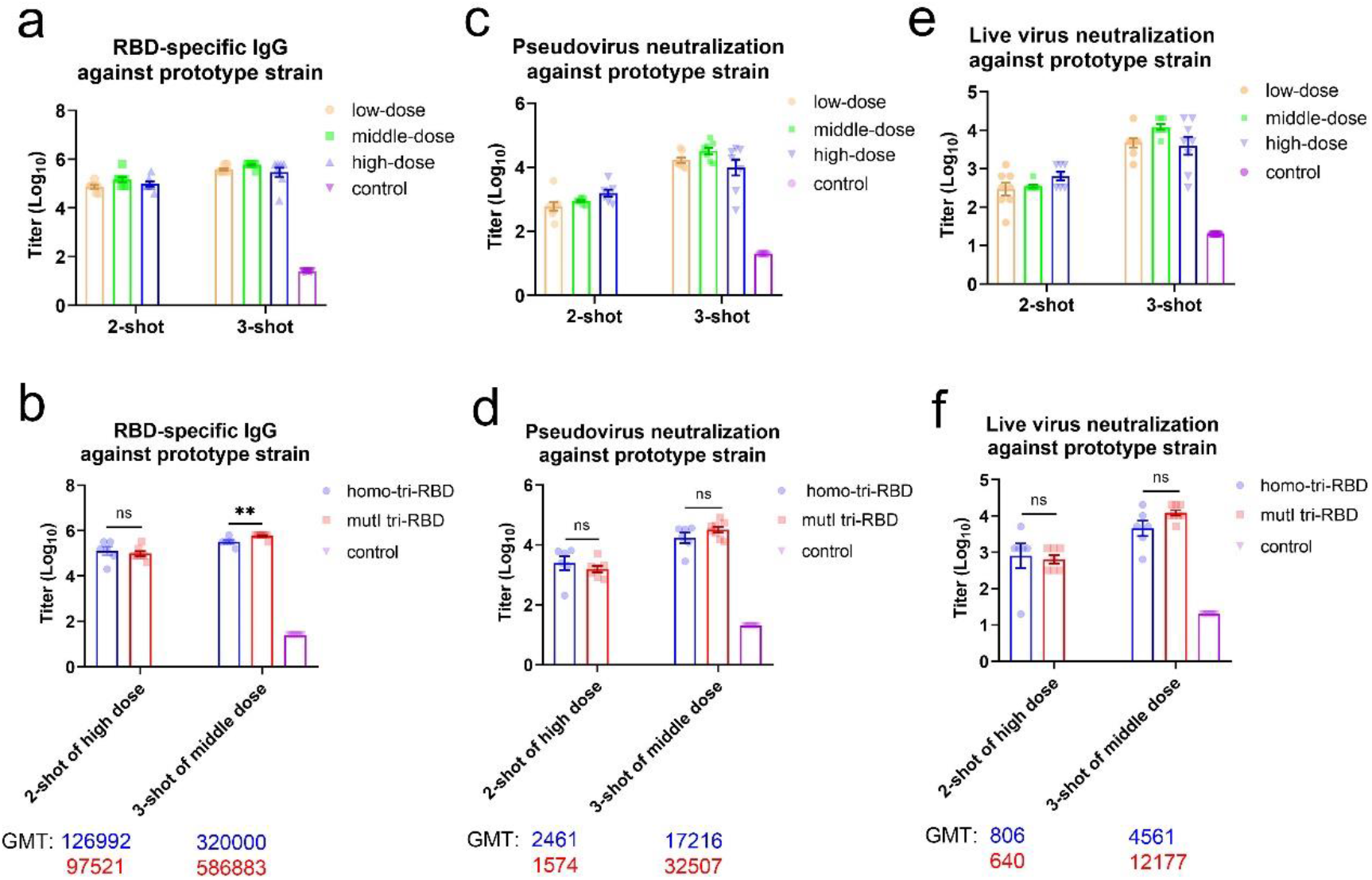
Immune responses against SARS-CoV-2 prototype strain elicited by mutI tri-RBD in mice compared with those induced by homo- tri-RBD. **a.**Dose-dependent responses of RBD-specific IgG against SARS-CoV-2 prototype strain elicited by mutI tri-RBD. The mice were immunized with two-shot or three-shot injections, and for each shot, three different doses were used including low (0.125 μg/dose), middle (0.5 μg/dose) and high (2.0 μg/dose) doses, respectively. The levels of IgG were detected with ELISA by using monomeric RBD of prototype strain. **b.**The levels of RBD-specific IgG against the prototype strain induced by mutI tri-RBD compared with those induced by homo-tir-RBD. **c.**Dose-dependent responses of neutralizing antibodies against SARS-CoV-2 pseudo-virus of the prototype strain elicited by mutI tri-RBD. The titers of neutralizing antibodies were assessed by using the pseudo-virus neutralization assays. **d.**The titers of neutralizing antibodies against the pseudo-virus of prototype strain induced by mutI tri-RBD in contrast with those induced by homo-tri-RBD. **e.**Dose-dependent responses of neutralizing antibodies against the live prototype SARS-CoV-2 virus elicited by mutI tri-RBD. The titers of neutralizing antibodies were assessed by using the live virus neutralization assays. **f.**The titers of neutralizing antibodies against the live virus of prototype strain induced by mutI tri-RBD in contrast with those induced by homo-tri-RBD. Data are presented as means±SEMs. P values were calculated with Student’s t-test. **P<0.01. ns: not significant.

**Fig. 3.**
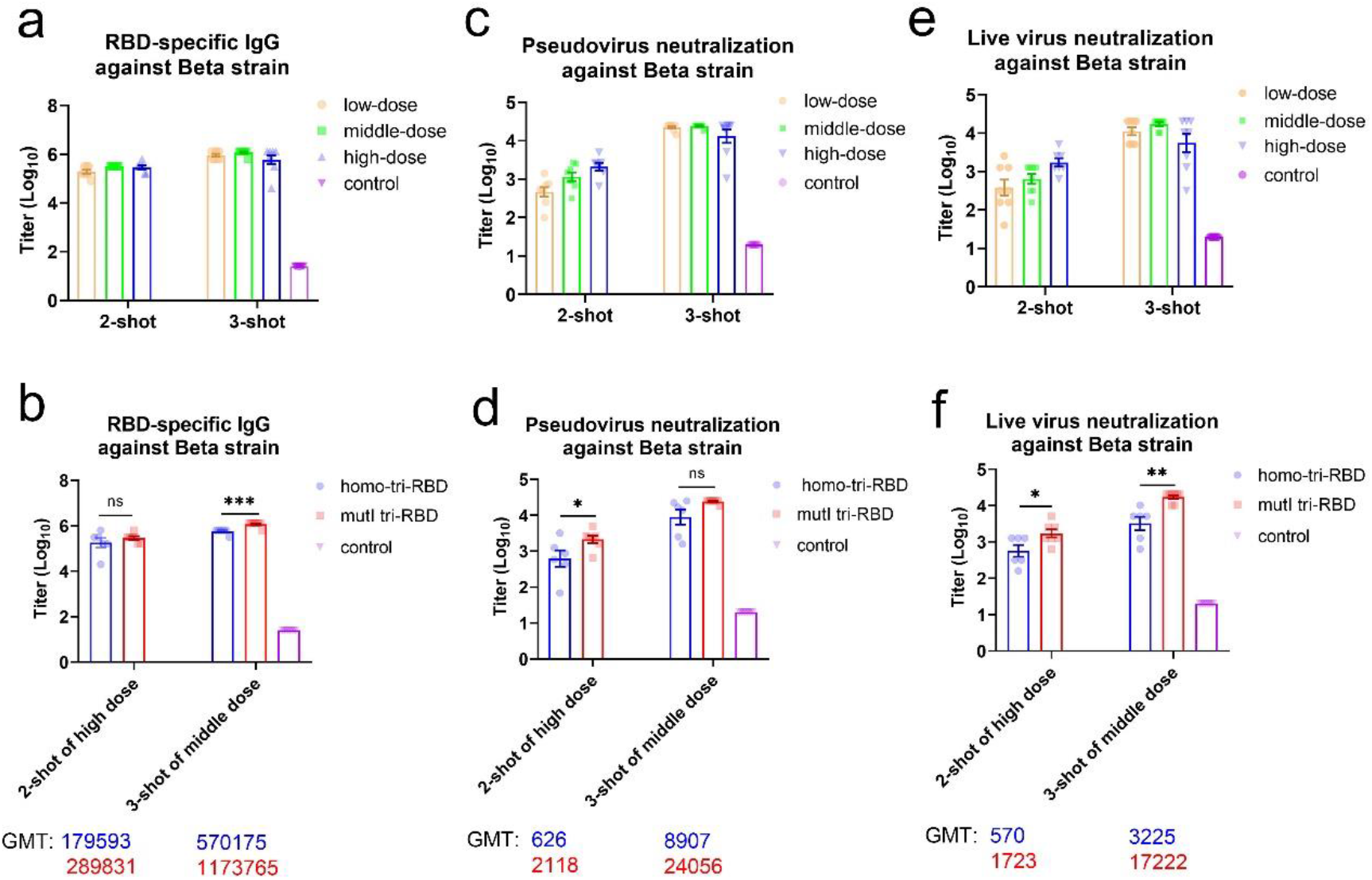
Immune responses against the Beta (B.1.351) strain of SARS-CoV-2 elicited by mutI tri-RBD in mice compared with those induced by homo-tri-RBD. **a.**Dose-dependent responses of RBD-specific IgG against the Beta (B.1.351) SARS-CoV-2 strain elicited by mutI tri-RBD. The levels of IgG were detected with ELISA by using monomeric RBD of Beta (B.1.351) strain. **b.**The levels of RBD-specific IgG against the Beta (B.1.351) strain induced by mutI tri-RBD compared with those induced by homo-tir-RBD. **c.**Dose-dependent responses of neutralizing antibodies against SARS-CoV-2 pseudo-virus of the Beta (B.1.351) strain elicited by mutI tri-RBD. The titers of neutralizing antibodies were assessed by using the pseudo-virus neutralization assays. **d.**The titers of neutralizing antibodies against the pseudo-virus of Beta (B.1.351) strain induced by mutI tri-RBD in contrast with those induced by homo-tri-RBD. **e.**Dose-dependent responses of neutralizing antibodies against the live Beta (B.1.351) SARS-CoV-2 virus strain elicited by mutI tri-RBD. The titers of neutralizing antibodies were assessed by using the live virus neutralization assays. **f.**The titers of neutralizing antibodies against the live virus of Beta (B.1.351) strain induced by mutI tri-RBD in contrast with those induced by homo-tri-RBD. Data are presented as means±SEMs. P values were calculated with Student’s t-test. *P<0.05, **P<0.01, ***P<0.001. ns: not significant.

Secondary structural composition analysis of the mutI tri-RBD protein by circular dichroism (CD) illustrated that the recombinant protein was well-folded, which consists of about 13.5% helix, 23.3% β strands, 11.4% turn and 51.8% others (Figure 1e and Extended Data Fig. 4). The sequence of one RBD contains 8 cysteine residues, and in the native structure of RBD in S protein, these cysteines form 4 intradomain disulfide bonds. Disulfide-linkages mapping by LC-MS showed that all these 4 disulfide bonds in RBD native structure also correctly formed in the recombinant mutI tri-RBD protein (Figure 1f and Extended Data Fig. 5). The secondary structure and disulfide bonds analyses suggest that each of the three RBDs in the recombinant mutI tri-RBD protein may be well-folded into its native structure. The stability of the protein was evaluated by differential scanning calorimetry (DSC) and the transition temperature (T_m_) was determined to be 47.2 ℃, indicating that the designed mutI tri-RBD protein was rather stable (Figure 1g).

**Fig. 4.**
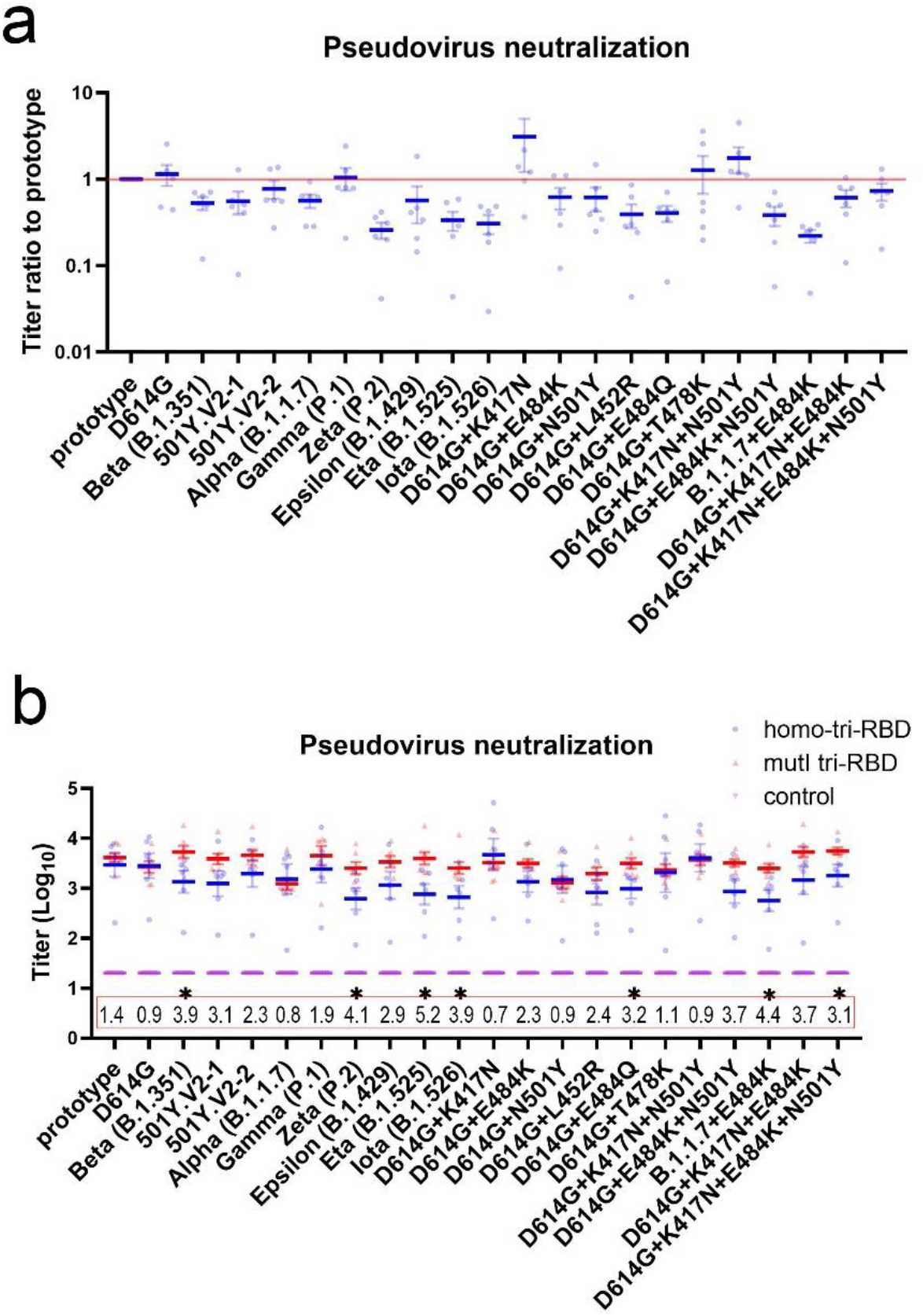
The neutralizing antibody responses against 22 various SARS-CoV-2 pseudo-virus strains elicited by mutI tri-RBD compared with those induced by homo-tri-RBD. **a.**Sensitivities to the neutralization of homo-tri-RBD immunized sera for the various pseudo-typed variants compared with that of the prototype strain. In this figure, GMT of the prototype strain is taken as reference, and GMT ratios between the variants and the prototype strain are displayed. Each serum sample is presented as a dot in the plot, and each serum sample was tested against all these variants. Data are presented as means±SEMs. **b.**Neutralizing antibody GMTs against the various pseudo-typed strains induced by mutI tri-RBD (red color) compared with those elicited by homo-tri-RBD (blue color). Numbers in the red box indicate the GMT ratios of mutI tri-RBD to homo-tri-RBD. P values were calculated with Student’s t-test. *P<0.05.

To characterize the bio-activity of the recombinant tri-RBD proteins, the binding capabilities of the designed mutI tri-RBD and homo-tri-RBD proteins with two anti-RBD monoclonal neutralizing antibodies (nAbs), i.e., MM43 and R117, were evaluated by using enzyme-linked immunosorbent assay (ELISA). MM43 can bind with not only the prototype RBD but also the Beta (B.1.351) and Kappa (B.1.617.1) RBDs. High binding activities with MM43 were observed both for the recombinant mutI tri-RBD and homo-tri-RBD proteins, indicating the formation of native conformation for individual RBD in the designed trimeric proteins (Figure 1h). R117 can only bind with the prototype and Kappa (B.1.617.1) RBDs but not with the Beta (B.1.351) RBD. ELISA measurements showed that owing to harboring the Beta (B.1.351) RBD, mutI tri-RBD exhibited relatively decreased binding activity with R117 than homo-tri-RBD, especially at lower protein concentrations (Figure 1h).

Then, the binding strength of the mutI tri-RBD protein with the receptor hACE2 was also quantified by surface plasmon resonance (SPR) assay. The dissociation constant KD was determined to be 3.20 × 10^−3^ nM, with the association rate constant k_a_ of 1.78 × 10^7^ *M*^−1^*S*^−1^ and the dissociation rate constant k_d_ of 5.67 × 10^−5^ *S*^−1^ (Figure 1i). SPR assay demonstrated that the designed mutI tri-RBD protein binds specifically to hACE2 with a high affinity, implying correct folding of the RBDs and high functionality of the recombinant mutI tri-RBD protein. These results supported the mutI tri-RBD to serve as an excellent immunogen.

### Both mutI tri-RBD and homo-tri-RBD elicited high level of immune responses against SARS-CoV-2 prototype strain in mice

BALB/c mice were immunized intraperitoneally with the recombinant mutI tri-RBD mixed with aluminum adjuvant, and the level of RBD-specific IgG as well as the titers of the pseudo- and live-virus neutralizing antibodies were measured to evaluate the immunogenicity of the designed mutI tri-RBD protein. In order to investigate the impacts of different immunization regimens and doses on the immune activities, the animals were immunized with two injections on day 0 and day 21, or with three vaccinations on day 0, day 7 and day 21, respectively, and for each injection three different doses, including low-dose (0.125 μg/dose), middle-dose (0.5 μg/dose) and high-dose (2.0 μg/dose), respectively, were applied. The sera of the immunized mice were collected on day 7 or day 14 after last vaccination. The level of RBD-specific IgG in the mice sera (collected on day 7 post immunization) was measured by using the enzyme linked immunosorbent assay (ELISA), in which the monomeric RBD from the SARS-CoV-2 prototype strain was used to coat the wells of the ELISA plates. An obvious dose-dependent response of the RBD-specific IgG was observed. Three-injection immunization regimen elicited distinctly higher immune response compared with the two-injection treatment, where the anti-RBD IgG titers were improved by 5.2, 4.0 and 3.0 times for the low-dose, middle-dose and high-dose vaccinations, respectively (Figure 2a). Both in the two-shot and three regimens, sera obtained from the mice immunized with low-dose (0.125μg/dose) antigen already exhibited high level of anti-RBD IgG compared with that of the mice in the control group, and middle-dose (0.5μg/dose) immunization improved the level of anti-RBD IgG. But, the high-dose vaccination did not further raise the geometric mean titer (GMT) of RBD-specific IgG (Figure 2a). Additionally, in order to compare the immunogenicity of the mutI tri-RBD with the homo-tri-RBD, BALB/c mice were also immunized with the homo-tri-RBD by two-injection of high dose (2.0 μg/dose) or three-injection of middle dose (0.5μg/dose). Notably, we observed that the RBD-specific IgG induced by the mutI tri-RBD was comparable to that elicited by the homo-tri-RBD in the two-injection immunization, and even obviously stronger in the three-injection immunization, which indicates similar, or even better, immunogenicity of the designed mutI tri-RBD compared with the homo-tri-RBD (Figure 2b).

Then, the GMT of the neutralizing antibodies against SARS-CoV-2 prototype strain induced by the mutI tri-RBD were assessed by using the pseudo- and live virus neutralization assays. The sera from the immunized mice (collected on day 7 post immunization) potently neutralized both the pseudo- and live virus infections, and similar to IgG response, a distinct dose-dependent response of the neutralizing antibodies was also observed. Three injections of the hybrid tri-RBD induced significantly higher neutralizing antibody titers than two shots, where the neutralizing GMTs against pseudo-virus were improved by 6.2~36.8 times for different doses, and also improved by 6.2~34.9 times against live SARS-CoV-2 prototype virus (Figure 2c,e). Moreover, in the two-injection regimen, higher dose per shot led to stronger neutralizing GMT against both the pseudo- and live virus. In the three-injection regimen, middle dose induced the highest neutralizing antibody titer, and the neutralizing GMT against live prototype virus reached as high as 12,177 (Figure 2e,f). More importantly, the neutralizing antibody response induced by the mutI tri-RBD was also compared with that of the homo-tri-RBD. In the two-shot immunization, the GMTs of the neutralizing antibodies induced by the mutI tri-RBD against both the pseudo- and live SARS-CoV-2 prototype strain were comparable to those elicited by the homo-tri-RBD, and interestingly in the three-shots vaccination, the neutralizing GMT values even tended to be higher for the mutI tri-RBD compared with the homo-tri-RBD although the difference was not statistically significant (Figure 2d,f). The results indicated that although only one of the three RBDs was derived from SARS-CoV-2 prototype strain, the neutralizing antibody responses against the prototype strain elicited by the mutI tri-RBD were not less, or even higher, than the homo-tri-RBD which contains three repeated RBDs from prototype strain.

It should be noted that owing to the good immunogenicity and protective efficacy in animals, the homo-tri-RBD has been approved by the China National Medical Products Administration (NMPA) to enter clinical trial (No. NCT04869592). Here, the immune responses both of IgG and neutralizing antibodies in mice suggested that the designed mutI tri-RBD exhibited similar or even better immunogenicity than the homo-tri-RBD against the prototype SARS-CoV-2 virus, although the former harbors only one-third prototype RBD in contrast to the latter. The results of animal experiments supported that the mutI tri-RBD may also serve as an effective vaccine candidate against the SARS-CoV-2 prototype strain.

### mutI tri-RBD induced significantly higher titer of neutralizing antibody responses against the Beta (B.1.351) variants than homo-tri-RBD

The primary purpose for the design of the mutI tri-RBD was to possibly develop a vaccine candidate with broadly neutralizing activities against SARS-CoV-2. We first tested whether the designed mutI tri-RBD can elicit cross-neutralizing antibody responses against one of the parent variants of the mutI antigen, i.e. the Beta (B.1.351). The level of specific IgG and neutralizing antibodies against Beta (B.1.351) variant induced by the mutI tri-RBD were evaluated in the mice sera (collected on day 7 post immunization) by using ELISA as well as pseudo- and live virus neutralization assays, which were then compared with those induced by the homo-tri-RBD.

The RBD-specific IgG detected by using the RBD monomer of the Beta (B.1.351) variant showed that the mutI tri-RBD can elicited strong antibody response both in two-shot and three-shot regimens, and the immune response was also dose-dependent. The level of specific IgG in the mice received three shots of the mutI tri-RBD was obviously improved than those received two shots. Both for two-shot and three-shot vaccinations, middle dose per shot caused slightly stronger IgG response than the low dose, but high dose did not further improve the specific IgG GMTs, suggesting possible immune saturation (Figure 3a). Notably, compared with the homo-tri-RBD, the designed mutI tri-RBD triggered the animal to produce significantly higher IgG titers in the three-shot immunizations, suggesting potentially better immunogenicity of the mutI tri-RBD against the Beta (B.1.351) variant (Figure 3b).

The neutralizing antibody response against the Beta (B.1.351) variant induced by the mutI tri-RBD also demonstrated a dose-effect relationship, where two injections with high dose and three injections with middle dose elicited the strongest neutralizing GMTs both in the pseudo- and live virus neutralization assays (Figure 3c,e). Especially, the GMT of neutralizing antibodies against the live virus of Beta (B.1.351) strain arrived at as high as 17,222 for three shots with middle dose of the mutI tri-RBD (Figure 3e). More importantly, compared with the homo-tri-RBD, the hybrid antigen elicited 3.4-fold and 2.7-fold higher neutralizing GMTs against the Beta (B.1.351) pseudo-virus in the two-shot and three-shot immunizations, respectively (Figure 3d). The live-virus neutralization assays further confirmed the pseudo-virus neutralization results, in which the neutralizing GMTs improved 3.0-fold and 5.3-fold, respectively, in the two-shot and three-shot vaccinations of the mutI tri-RBD than those of the homo-tri-RBD (Figure 3f). Therefore, regarding to the Beta (B.1.351) variant, the mutI tri-RBD exhibited superior immune efficacy than the homo-tri-RBD. The above results indicated that the mutI tri-RBD not only maintained the strong neutralizing antibody responses against the prototype SARS-CoV-2 strain, but also induced significantly improved neutralizing antibody titers against the Beta (B.1.351) variant in contrast to the homo-tri-RBD.

The above results demonstrated that the designed mutI tri-RBD, which harbored the immunogens from three typical SARS-CoV-2 strains, elicited robust cross-variant neutralizing activities both against the prototype strain and the so called “immune escape” variant, i.e., the Beta (B.1.351) strain. Especially, the neutralizing titers against the Beta (B.1.351) variant were significantly improved for the mutI tri-RBD in contrast with the homo-tri-RBD.

### mutI tri-RBD elicited broadly neutralizing activities against SARS-CoV-2 compared with homo-tri-RBD

Besides the prototype and Beta (B.1.351) SARS-CoV-2 strains, we also investigated whether the designed mutI tri-RBD can also elicit broad neutralizing antibody responses against other currently-circulating or possibly-emerging SARS-CoV-2 variants. The pseudo-viruses of ten other circulating strains, including D614G, 501Y.V2-1, 501Y.V2-2, Alpha (B.1.1.7), Gamma (P.1), Zeta (P.2), Epsilon (B.1.429B), Eta (B.1.525) and Iota (B.1.526), as well as eleven corresponding single and combinatorial mutations in RBD were selected^24,25^, and the neutralizing antibody GMTs against these different SARS-CoV-2 variants were tested by using pseudo-virus neutralization assays. The sera collected on day 14 post-vaccination from the mice immunized with two shots of high dose were used for all the pseudo-virus neutralization reaction examinations. The prototype and Beta (B.1.351) strains were also re-tested by using the 14-day post immunization sera as comparison. In order to assess whether the mutI tri-RBD has superior performances in the broad neutralization against these various SARS-CoV-2 strains than the homo-tri-RBD, the GMT values elicited by the mutI tri-RBD were compared with those induced by the homo-one.

We first examined the neutralization reactions of the homo-tri-RBD immunized sera with the selected 21 SARS-CoV-2 variants to determine whether these variants exhibited resistances to the antibody neutralization elicited by homo-tri-RBD. Although homo-tri-RBD can induce neutralizing antibodies against all these tested variants in comparison with the control group, some variants also displayed obviously reduced sensitivities to the neutralization of the immunized sera compared with the prototype strain. Among the 10 circulating variants, Beta (B.1.351), 501Y.V2-1, 501Y.V2-2, Alpha (B.1.1.7), Zeta (P.2), Epsilon (B.1.429), Eta (B.1.525) and Iota (B.1.526), exhibited decreased neutralization susceptibilities to varying extents (Figure 4a). Notably, except Alpha (B.1.1.7), these variants all harbored E484K or L452R mutations, suggesting the importance of these two residues in immune resistance. This point was also confirmed by the corresponding single and combinatorial mutations, where distinctly reduced neutralization sensitivities were observed for the E484K- or L452R-carrying mutations (Figure 4a). In addition, the single mutations of E484Q or N501Y also displayed obviously reduced neutralization sensitivities to the antibodies elicited by homo-tri-RBD. However, the neutralization reactions against the K417N and T478K single-mutations were improved, implying that these two residues do not lead to immune resistance to tri-RBD (Figure 4a). This is consistent with the experimental results of Zhang, et al., and they found that K417N mutation could lead to enhanced neutralization activity^24^.

Then, we tested whether the designed mutI tri-RBD could induce more robust neutralization activities than tri-RBD against these 21 SARS-CoV-2 variants, especially the homo-tri-RBD-resistant variants discussed above. The results showed that for almost all these tested strains, the neutralization GMTs induced by mutI tri-RBD were comparable to or significantly higher than those induced by tri-RBD. For all the E484K- or L452R-harboring variants of the 10 circulating variants, including Beta (B.1.351), 501Y.V2-1, 501Y.V2-2, Gamma (P.1), Zeta (P.2), Epsilon (B.1.429), Eta (B.1.525) and Iota (B.1.526), the neutralization activities induced by mutI tri-RBD were significantly improved by 1.9 to 5.2-folds in contrast to those induced by tri-RBD (Figure 4b). Regarding to the E484K, L452R, as well as E484Q single-mutations, the neutralization sensitivities were raised 2.3, 2.4, and 3.2-folds, respectively, elicited by the mutI tri-RBD (Figure 4b). Most importantly, as discussed above, the E484K, L452R and E484Q mutations could cause obvious resistances to the sera immunized with homo-tri-RBD, which may partly reduce the efficacy of the homo-tri-RBD candidate vaccine. However, encouragingly, the mutI tri-RBD enables a substantial increase of the neutralizing titers against these homo-tri-RBD-resistant variants, indicating potently broad neutralizing capabilities induced by mutI tri-RBD (Figure 4b). E484K has been believed to be the most important “immune escape” mutation, and our pseudo-virus neutralization assays displayed that the mutI tri-RBD also induced robustly higher neutralizing antibody responses against the E484K-carrying combinatorial mutations in contrast to homo-tri-RBD. It should be mentioned that for D614G, B.1.1.7, D614G+K417N, D614G+N501Y, D614G+K417N+N501Y and D614G+T478K variants, the mutI tri-RBD elicited similar neutralizing antibody response compared with homo-tri-RBD (Figure 4b).

It should be mentioned that Kappa (B.1.617.1) variant was another parent strain for the construction of mutI tri-RBD. At present, the pseudo- and live viruses of Kappa (B.1.617) variant are not yet obtained by our group, and thus the neutralizing activities against Kappa (B.1.617.1) variant elicited by the mutI tri-RBD could not be directly detected. Nevertheless, the neutralizing activities against each of the single mutations harbored by the Kappa (B.1.617.1) and Delta (B.1.617.2) variants, including L452R, E484Q and T478K, were measured in this study. The T478K mutation did not lead to immune resistance (Figure 4a), and for L452R and E484Q mutations, mutI tri-RBD induced significantly stronger neutralizing GMTs than homo-tri-RBD (Figure 4b). Therefore, we speculate that the mutI tri-RBD may potentially induce superior neutralizing activities against the Kappa (B.1.617.1) and Delta (B.1.617.2) variants compared with homo-tri-RBD.

Zahradník et al. revealed that the E484K, L452R and N501Y mutations exhibited the strongest convergent evolution in RBD, which could be acquired independently across different lineages during the evolution of the virus^11^. These mutations were considered to be of adaptive advantage, suggesting the possible recurrence of them individually or combinedly in the future. The designed mutI tri-RBD may also induce neutralizing antibody response potentially against the future-emerging SARS-CoV-2 variants.

Taken together, the designed mutI tri-RBD candidate vaccine could not only induce robust neutralizing activities against the prototype and the parent (B.1.351) SARS-CoV-2 strains, but also have the potential to provide broadly neutralizing capabilities against many other currently-circulating and even future-emerging variants. It is important to point out that in our previous study (not published), homo-tri-RBD could fully protect the transgenic mice against the live SARS-CoV-2 virus challenges (Extended Data Fig. 6). The more robust neutralizing antibody GMTs induced by the mutI tri-RBD may indicate its more broadly protective effectiveness compared with homo-tri-RBD.

## DISCUSSION

SARS-CoV-2 is continually evolving to acquire mutations that may impact viral infectivity, transmissibility and antigenicity. Alarmingly, growing evidences have demonstrated that some emerging variants are partly resistant to the neutralizing antibodies induced by natural infection or vaccination, which has raised serious concerns about the efficacy of the existing vaccines. It is urgent to develop broadly protective vaccines to combat with the currently circulating, or even future occurring, immune-resistant SARS-CoV-2 variants. The RBD in S protein mediates infection of the virus to host cell, which is the principal target for vaccine design. The naturally trimeric arrangement of RBDs in S protein, as well as its unique structural characteristics, allows the realization of a mutI tri-RBD as a broadly protective vaccine candidate, in which the antigens derived from various SARS-CoV-2 circulating strains were integrated into a single immunogen. The designed mutI tri-RBD not only induced strong neutralizing antibody responses against the prototype SARS-CoV-2 strain, but also elicited robustly broad neutralizations against the so called “immune escape” variants. Owing to harboring the immune-resistant mutations as well as the evolutionarily convergent hotspots, the mutI tri-RBD may also have the potential to provide neutralizing capability against the future-emerging SARS-CoV-2 variants. Compared with the homo-tri-RBD, the hybrid immunogen displayed superior broad-neutralization performances against SARS-CoV-2.

The vaccines specifically targeted at the Beta (B.1.351) SARS-CoV-2 variant has recently begun to be developed, and the anti-Beta (B.1.351) vaccine candidate has been tested in combination with the prototype as a bivalent vaccine or as a heterologous prime boost^26,27^. Here, we provided another novel strategy for the construction of a vaccine with broad neutralization capability, which incorporated different variant-specific antigens into a single hybrid immunogen. Compared with the conventional multivalent strategy, our method has the advantages of lower time costs and higher production benefits. In the mutI tri-RBD vaccine candidate, the aluminum adjuvant was adopted, which can improve the accessibility and affordability of the vaccine. All these features prospectively support the further development of the mutI tri-RBD vaccine candidate in clinical trials.

## METHODS

### Structural modelling and molecular dynamics simulations of the designed mutI tri-RBD and homo-tri-RBD proteins

We aim to construct a mutation-integrated trimeric form of RBD (mutI tri-RBD) to possibly mimic the trimeric RBD arrangements in the natural S protein and integrate different immunodominant antigens into a single molecule. In mutI tri-RBD, the RBD region (residues 319-537) was truncated from the S proteins of the prototype, Beta (B.1.351) and Kappa (B.1.617.1) SARS-CoV-2 strains, respectively, and then these three RBDs were connected end-to-end by using their own long loops at the N- and C-termini of the truncated regions without introducing exogenous linker. As comparison, the homo-tri-RBD was also constructed in which the three RBDs were all truncated from the prototype strain. The possible three-dimensional structures of the designed mutI tri-RBD and homo-tri-RBD proteins were modelled with Modeller9.23 software^22^ by using the native structure of S trimer (PDB accession code 6zgi for homo-tri-RBD, as well as 6zgi and 7lyl for mutI tri-RBD) as the template. A total of 10 structures were generated both for mutI tri-RBD and homo-tri-RBD, and the structure with the lowest value of the DOPE assessment score^28^ was picked as the best model.

In order to assess the stability as well as the intrinsic dynamics of the modelled structures, a 200ns atomic MD simulation was performed both for the mutI tri-RBD and homo-tri-RBD. All the MD simulations were carried out by using Gromacs 2019 with the Charmm27 force field. The trimeric structure generated by Modeller was solvated using TIP3P water molecules in a cubic box, with the protein atoms being at least 1.6 nm away from the box edges. A total of 24 and 21 CL-ions were added into the water box to neutralize the net charges of the simulation systems for the mutI tri-RBD and homo-tri-RBD, respectively. The prepared system was then subjected to energy minimization with the steepest descent algorithm to ensure that the maximum force in the system was below 1000.0 kJ/mol/nm. After energy minimization, a 100 ps NVT simulation with position restraints on protein atoms was performed at 300K, which was followed by a 100 ps NPT simulation with position restraints. Finally, the production simulation without any position restraint was run for 200 ns and the snapshots were collected every 10 ps to obtain the simulation trajectory. In the simulation, a time step of 2 fs was used and all h-bonds were constrained with LINCS algorithm. Both in the mutI tri-RBD and homo-tri-RBD, each RBD unit contains four disulfide bonds, and these disulfide bonds were included in the construction of the topology files for the simulation systems. A cutoff value of 1.0 nm was used for the calculation of both short-range electrostatic and van der Waals interactions. Long-range electrostatic interactions were computed by using the Particle-Mesh Ewald (PME) algorithm. During the simulation, the temperature and the pressure of the system were maintained at 300K and 1 bar with the velocity rescaling and the isotropic Parrinello-Rahman coupling methods, respectively. Based on the MD simulation trajectories, the changes in the root mean square deviation (RMSD) of the C_α_ atoms of the system as a function of time were calculated by using the built-in tool of “gmx rms” in Gromacs to evaluate the stability of the trimeric RBD proteins, and the motion movies were generated by using Chimera software to display the intrinsic conformational movements of the trimeric RBD proteins.

### Expression and purification of the designed recombinant mutI tri-RBD and homo-tri-RBD proteins

The gene sequences of the designed mutI tri-RBD and homo-tri-RBD proteins were codon optimized for the transient expression in the mammalian cells. In the designed trimeric schemes, mutI tri-RBD is composed of three RBD regions derived from the S proteins of the prototype, Beta and Kappa SARS-CoV-2 strains, respectively, connected end-to-end with each other. Homo-tri-RBD consists of three copies of the RBD region from the prototype strain. For gene cloning of these designed proteins, signal peptide and Kozak sequences were added to the N-terminal of the expressed protein sequences, which were then inserted into the PTT5 plasmid via the *Hin* dIII and *Not* I restriction sites to construct the recombinant plasmids for the expression of the trimeric RBDs in the mammalian HEK293T cells. The sequences of the constructed plasmids for the mutI tri-RBD and homo-tri-RBD proteins were verified by gene sequencing. Then, the generated plasmids were transfected into the HEK293T cells for transient expression. After 3-5 days culture, the culture supernatants from the transfected cells were harvested and purified by chromatography. During the chromatographic purifications, the isolated proteins from the eluted peaks was analyzed with SDS-PAGE. Then, the recovered protein sample was finally purified by ultrafiltration with the membranes of 30-kDa molecular weight cutoff. The purity of the produced proteins was determined by the size exclusion-high-performance liquid chromatography (SEC-HPLC) using TSKgel®G2500PW column.

### MALDI-TOF-MS analysis to measure the molecular weight

MALDI-TOF-MS (Matrix-assisted laser desorption ionization-time of flight mass spectrometry) analysis was carried out with AB Sciex 4800 Plus MALDI TOF analyzer to measure the molecular weight of the recombinant mutI tri-RBD^29,30^. The protein sample was exchanged into water to the concentration of 5 mg/mL, and 0.5 μL of the sample was spotted onto the MALDI target plate, followed by air drying at room temperature. The MALDI plate was then covered with 0.5 μL of 0.5 mg/mL sinapinic acid (SA) dissolved in 0.1% TFA and 50% CAN, and also dried by air at room temperature. The MALDI TOF/TOF analyzer was equipped with a Nd:YAG 355nm laser. Parameters were set as follows: Linear mode; repetition rate laser: 200 Hz; mass range: 20000-150000 Da; 400 shots accumulated per profile.

### UPLC-MS analysis to identify the N- and O-glycosylation sites

To identify the N-glycosylation sites on the protein by using UPLC-MS (ultra-high performance liquid chromatography coupled with mass spectrometry), the asparagine (N)-liked glycans were firstly cleaved by PNGase F endoglycosidase in ^18^O-water at 37 ℃ for 18h, to label the N-glycosylation site with ^18^O. Then, the N-Deglycosylated protein was denatured and reduced with denaturant and DTT at 37 ℃ for 30 min. Subsequently, the protein was alkylated with iodoacetamide (IAA) in the dark at room temperature for 30 min, which was then digested with Glu-C enzyme at 37 ℃ for 18 h. Then the digested peptides were separated by ultra performance liquid chromatography (UPLC) using a Waters ACQUITY UPLC Peptide BEH C18 column (130Å, 1.7μ*m*, 2.1 *mm* × 150*mm*) at the temperature of 60 ℃, with a flow rate of 0.3 mL/min. The eluting peptides were analyzed on the Q Exactive Plus mass spectrometer (MS) in positive ion mode. The MS spectra were acquired with a scan range of 300-2000 m/z and a resolution of 35000. The LC/MS raw files were processed with BioPharma Finder software (version 3.2) to identify the N-glycosylation sites.

To identify the O-glycosylation sites, the protein sample was firstly denatured with urea and reduced with DTT at 37 ℃ for 30 min, followed by the alkylation with NEM at 37 ℃ for 2h. The sample was pre-cleaved with Lys-C enzyme at 37 ℃ for 4h. Then, the SialEXO (40 U/μL, Genovis/G1-SM1-020), OpeRATOR(40 U/μL,Genovis/G2-OP1-020), PNGase F (NEB/P0704L) and 20mM Tris buffer (pH=7.5) were added into the protein sample, in which the protein was digested into O-glycopeptides at 37 ℃ overnight. The resulted peptides were analyzed by using reverse-phase UPLC coupled with QE to identify larger O-glycosylated peptides. Smaller peptides in the digestion product were purified using the Graphite Spin column (Pierce™, cat. 88302) according to manufacturers’ instructions. After that, the O-glycopeptides were analyzed by HILIC (hydrophilic interaction chromatography) UPLC coupled with QE. And the raw data files were analyzed with BioPharma Finder software to identify the O-glycosylation sites and potential O-glycan forms.

### UPLC-MS analysis to map the disulfide bond locations

To identify the disulfide bond locations in the recombinant mutI tri-RBD protein by using the UPLC-MS peptide mapping method, both reduced and non-reduced protein samples were prepared. In sample preparation, the N-ethylmaleimide (NEM) solution was firstly added into the sample to block free sulfhydryl groups in the protein at room temperature for 2 h. Then, the protein was transferred into a digestion buffer and pre-cleaved with Lys-C at 37 ℃ for 4 h followed by an additional digestion with Glu-C at 37 ℃ overnight. Half of the digested sample was taken as the non-reduced sample, and the other half was further subjected to reduction and alkylation to prepare the reduced sample. In reduction and alkylation reactions, the digested peptides were reduced with DTT at 37 ℃ for 30 min and alkylated with IAA in the dark at room temperature for an additional 30 min. Then the resulted peptides were separated both for the reduced and non-reduced samples by UPLC using a C18 column (130Å, 1.7μ*m*, 2.1 *mm* × 150*mm*) with the column temperature of 60 ℃, and the reversed-phase chromatography of the sample was performed for 95 min with a flow rate of 0.3 ml/min and a UV detection wavelength of 214 nm. Subsequently, the separated peptides were detected and analyzed by Q Exactive Plus MS in positive ion mode, with m/z range of 300-2000 and the resolution (Full MS/MS2) of 35000/17500. The MS raw data files were processed using BioPharma Finder software (version 3.2). The presence of disulfide-bonds containing peptides were determined through comparative analysis of the reduced and non-reduced peptide samples, and were shown as indicated in the chromatogram as shaded peaks. The free sulfhydryl of a certain cysteine was quantified through the ratio of MS area of the NEM modification containing peptides against MS area of all peptides in the reduce sample.

### CD spectroscopy to estimate the secondary structural composition

CD spectroscopy was performed on Applied Photophysics following the operation procedure provided by the manufacturer. The protein sample was dialyzed into the buffer composed of 5 mM PB, and then placed to the spectrophotometer cell. Both far-ultraviolet (UV) and near-UV spectra were acquired in the wavelength range of 190-250 nm and 250-3040 nm, respectively, with spectral resolution of 0.5 nm and band-width of 1.0 nm. All CD spectra were obtained with 0.5 S per point at 25 ℃. Both the spectra of the sample and the buffers under the same experimental conditions were recorded, and the actual protein spectra were obtained by subtracting the spectra of the buffers. The scan measurement was repeated 6 times and the average result was calculated to ensure the reliability of the measurement. The acquired CD data were analyzed with the BeStSel Server (http://bestsel.elte.hu/index.php)^31^ to estimate the content of different secondary structures in the protein.

### DSC analysis to characterize protein stability

The protein sample was prepared at the concentration of 2.0 mg/mL in origin formulation buffer and 0.3 ml of the sample was used to perform the Differential scanning calorimetry (DSC) experiment with the TA Instrument Nano-DSC. The thermograms were obtained over the temperature range from 5 ℃ to 95 ℃ using a scan rate of 1 ℃/min. The experimental data were processed by Launch NanoAnalyze software supplied with the DSC instrument.

### SPR measurement to detect the binding affinity with hACE2

The surface plasmon resonance (SPR) measurements were carried out by Biakore 8K (GE Healthcare) with NTA chips. The His-tagged receptor protein hACE2 was dissolved in HBS-T buffer (HBS buffer and 0.05% Tween20) and immobilized onto the NTA chip. The protein sample was diluted with HBS-T buffer at the concentrations of 0.0173, 0.0346, 0.0692, 0.1385, and 0.277 μg/ml, which were then flowed over the chip surface at a rate of 30 μL/min for 120 s. The bound protein was then dissociated for an additional 120s. During the association and dissociation processes, the real-time response SPR signal was recorded. After that, the sensor chip was regenerated using 350 mM EDTA regeneration solution for 120s with a flow rate of 30 μL/min. The recorded data were processed using BiacoreTM Insight Evaluation software and the binding kinetics were analyzed with a 1:1 binding model to obtain the association rate constant k_a_, the dissociation rate constant k_d_, and the dissociation constant KD.

### ELISA to evaluate the binding activities with various monoclonal neutralizing antibodies

The binding activities of the recombinant mutI tri-RBD and homo-tri-RBD proteins with two monoclonal neutralizing antibodies (nAbs), including MM43 and R117 (Sino Biological Inc., China. Cat: 40591-MM43 and 40592-R117), were evaluated by using enzyme linked immunosorbent assay (ELISA). To detect the binding activity with the anti-RBD monoclonal nAbs, the protein samples of mutI tri-RBD and homo-tri-RBD were prepared at the starting concentration of 1.0 μg/mL, which were then subjected to 2-fold serial dilutions and coated on the 96-well microplate with 100 μL per well at 2-8 ℃ overnight. Subsequently, the plate was washed 3 times with phosphate-buffered saline (PBS) containing 0.1% Tween 20 (PBST) and then blocked with 100 μL blocking buffer per well, followed by incubation at 37 ℃ for 2 h. After washing the plate 3 times with PBST, the nAb sample was diluted to 1.0 μg/mL and added into the wells with 100 μL per well, which was then incubated at 37 ℃ for 1 h. After that, the plate was washed 3 times with PBST and the HRP-conjugated secondary antibody (goat anti-mouse IgG) at 1:10000 dilution was added into the wells of the plate. The plate was again incubated at 37 ℃ for 1 hour and washed 3 times with PBST. Subsequently, 50 μL tetramethylbenzidine (TMB) and 50 μL hydrogen peroxide solutions were added to start the color reaction. After color development for 5 min, the reaction was stopped using 0.2 M sulfuric acidic solution with 50 μL per well, and absorbance at 450 nm was read by plate reader. The background absorbance at 630 nm was also measured, and then the difference between absorbance at 450 nm and 630 nm (OD_450/630nm_) was obtained to detect the specific binding of the mutI tri-RBD and homo-tri-RBD proteins with the nAbs.

### Mouse immunization protocols

Female BALB/c mice with 6-8 weeks old were used in this study, which were purchased from Beijing Vital River Laboratory Animal Technology Co., Ltd., China. All the animal experimental procedures were carried out under the approvement of the Institutional Animal Care and Use Committee of the National Vaccine and Serum Institute of China.

To evaluate the impacts of different immunization regimens as well as vaccination doses on the immune responses, a total of 6 groups of mice (8 mice in each group) were intraperitoneally vaccinated with the mutI tri-RBD vaccine candidate mixed with aluminum adjuvant. In these experimental groups, the mice were immunized with two shots on day 0 and day 21, or with three shots on day 0, day 7 and day 21, respectively. For each shot, three different doses, including low-dose (0.125 μg/dose), middle-dose (0.5 μg/dose) and high-dose (2.0 μg/dose), respectively, were used. Another 8 mice were injected with only adjuvant on Day 0, Day 7 and Day 21, serving as the control group. In order to compare the immunogenicity of the mutI tri-RBD with the homo-tri-RBD, another two groups of BALB/c mice (6 mice in each group) were also immunized with the homo-tri-RBD plus aluminum adjuvant by two-shot of high dose (2.0 μg/dose) or three-shot of middle dose (0.5μg/dose). For all these groups, the sera of the immunized mice were collected at 7 days and 14 days after last immunization.

### ELISA to measure the titer of specific IgG in the sera of the immunized mice

The titer of the RBD-specific IgG in the serum of the immunized mice were measured with enzyme linked immunosorbent assay (ELISA) by using monomeric His-tagged RBDs of the prototype and Beta (B.1.351) SARS-CoV-2 strains, purchased from Sino Biological Inc., China (Cat: 40592-V08B and 40592-V08H85). In ELISA, the His-tagged RBD, was diluted with coating buffer to the concentration of 1 μg/mL, which was coated on the 96-well plate with 100 μL per well at 2-8 ℃ overnight. After washing 3 times with phosphate-buffered saline (PBS) containing 0.1% Tween 20 (PBST), the plate was blocked using blocking buffer with 100 μL per well and incubated at 37 ℃ for 2 h. Then, the plate was washed 3 times with PBST, and the serum samples were diluted by 2-fold serial dilutions and added into the well with 100 μL per well. After incubating the plate at 37 ℃ for 1 h and washing 3 times with PBST, the HRP-conjugated secondary antibody (goat anti-mouse IgG) at 1:10000 dilution was added into the wells of the plate. The plate was again incubated at 37 ℃ for 1 hour and washed 3 times with PBST. Subsequently, color reaction was performed for 10 min by adding the color development solution, which was then stopped using sulfuric acidic solution. Both absorbance at 450 nm and 630 nm were measured and the difference between them, i.e., OD_450/630nm_, was obtained. To evaluate the titer of RBD-specific antibodies, the OD_450/630nm_ value of the well without adding serum sample was taken as the blank control, and 2.1 times of the control value was used as the cutoff for positive results. The titer of RBD-specific IgG was determined as the reciprocal of the maximum dilution of the serum, where the OD_450/630nm_ value is equal to or greater than the cutoff value.

### SARS-CoV-2 pseudo-virus neutralization assays to evaluate the neutralizing antibody titer

A total of 22 SARS-CoV-2 pseudo-viruses were used to evaluate the neutralizing antibody titers in the sera from immunized mice. To evaluate the neutralization activities of the sera against these pseudo-viruses, the serum sample was serially diluted by 5-fold at the starting dilution of 1:40 with the cell culture medium (DMEM containing 10% Fetal Bovine Serum, 25 mM HEPES and 1% penicillin streptomycin). The serially diluted serum samples were added into the well of the plate with 50 μL per well. The SARS-CoV-2 pseudo-virus was diluted 9 times with the cell culture medium to the titer of 1.3×10^4^ TCID_50_ per mL and added into the well using 50 μL per well to mix with the serum, which was then incubated at 37 ℃ for 1 h. 50 μL per well medium was also mixed with the pseudo-virus as control. 100 μL per well medium served as blank. The Huh-7 cells were digested and suspended in the culture medium with viable cell density of 2×10^5^ per mL, which were then added into the well with 100 μL per well. After culture with 5% CO_2_ at 37 ℃ for 20 ~ 24 h, the culture medium was removed, and the cells were lysed and the luciferase activity was evaluated as the relative light unit (RLU) value. The neutralizing antibody titer was determined as the reciprocal of the dilution of the serum for 50% neutralization of viral infection, which was calculated using the Reed-Muench method.

### Live SARS-CoV-2 virus neutralization assays to evaluate the neutralizing antibody titer

To evaluate the neutralization activity against the live SARS-CoV-2 virus, the live virus neutralization assay was performed in a BSL3 laboratory of Guangdong Provincial Center for Disease Control and Prevention. Both the prototype (2020XN4276 strain, isolated by Guangdong CDC, China, GISAID: EPI_ISL_413859) and Beta (B.1.351) (20SF18530 strain, isolated by Guangdong CDC, China, GISAID: EPI_ISL_2536954) viruses were used in the live virus neutralization assays. The mouse serum was serially diluted and mixed with an equal volume of 100 TCID_50_ live SARS-CoV-2 virus. After incubation at 37 ℃ for 2 h, the serum-and-virus mixed solution was added into the well of the plate containing Vero-E6 cells with density of 2×10 ^5^ per mL, which was then cultured at 37 ℃ for 5 to 7 days. Both cell and virus controls were also set up as comparison. Subsequently, the inhibition of virus infection to the cells was observed, and the neutralizing antibody titer against the live SARS-CoV-2 virus was measured as the reciprocal of the serum dilution for 50% neutralization of viral infection.

### Statistical analysis

Comparisons for mutI tri-RBD and homo-tri-RBD were carried out by using unpaired Student’s t-tests. *P<0.05, **P<0.01, ***P<0.001, ****P<0.0001, ns: not significant.

## Acknowledgements

We thank Dr. Zhi Hua Xiao for assisting with protein expression. We also thank Prof. Hui Wang for providing us the prototype SARS-CoV-2 virus in mouse challenge experiments. We are grateful to Dr. Qi Zhang for assistance with protein physico-chemical characterization.

## Author contributions

Q. M. Li and J. Zhang conceived and supervised the project. Q. M. Li, Yu Liang, J. Zhang designed the experiments. J. G. Su conducted structure modelling and simulations. H. Zhang and F. J. Shen performed cell culture and protein expression. Y. Q. Jin, J. W. Hou, Z. M. Liu and Y. N. Hou conducted protein purification. X. F. Zhang, L. F. Du and S. Shao performed protein structural characterization. X. F. Zhang and J. J. Wu performed vaccine formulation and mice immunization. Z. Y. Hu, P. He verified the experimental methods. L. F. Du, F. Tang, Z. J. Ma, N. Liu and S. Y. Ren performed ELISA tests. W. J. Huang, M. Y. Wang, Z. B. Han, Z. H. Lei, S. Chen, F. Zheng and S. Liu constructed pseudotyped virus and performed neutralization assay. C. W. Ke and R. Y. Yuan conducted live virus neutralization tests. G. Z. Wu, Y. Liang, X. Zheng, N. Liu, W. M. Zhou, W. Zhen performed the mouse challenge experiments. Q. M. Li, J. Zhang, Y. Liang and J. G. Su analyzed the experimental data. J. G. Su, Z. B. Han, Y. Liang wrote the manuscript. Q. M. Li and J. Zhang revised the manuscript.

## Supporting Materials

**Extended Data Fig. 1.**
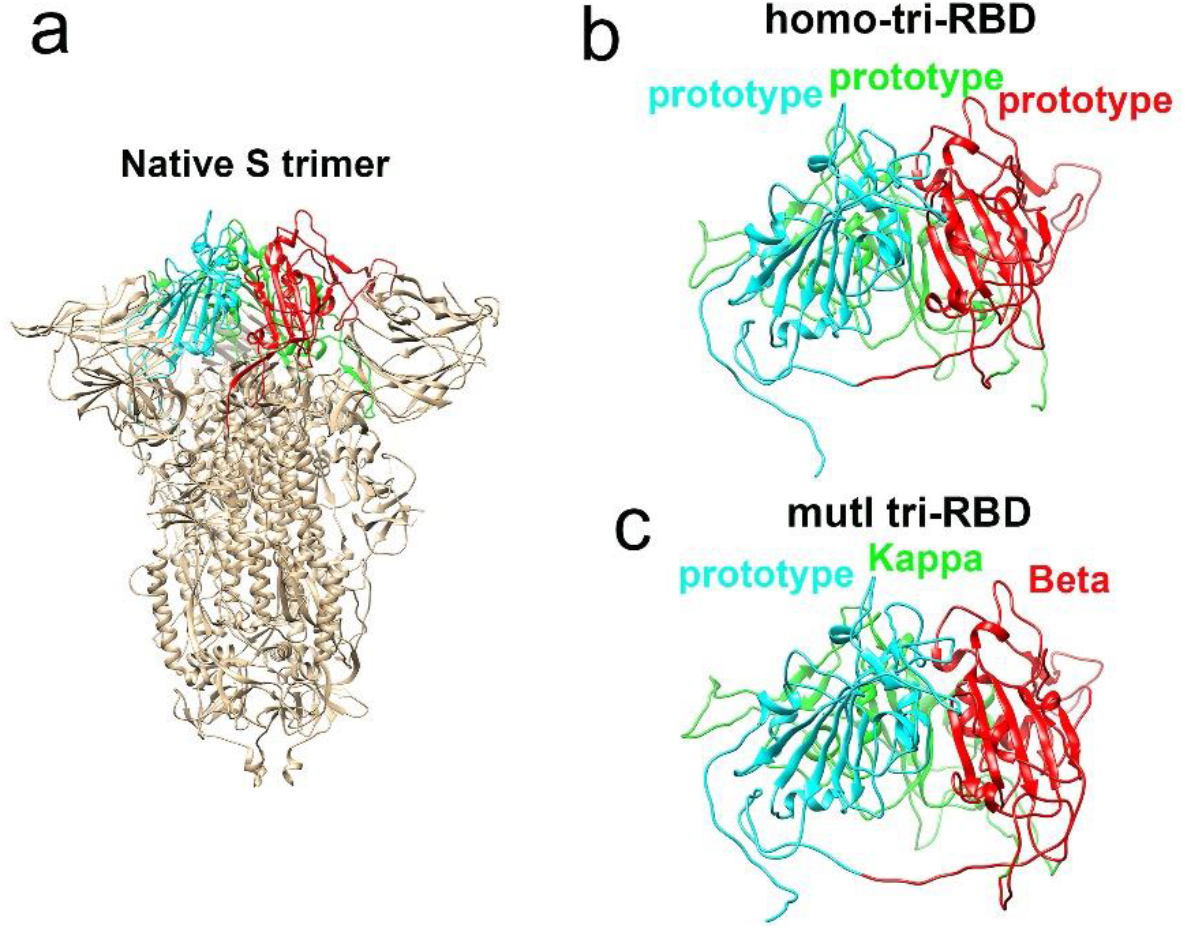
Structural modelling and molecular dynamics (MD) simulation of the designed mutI tri-RBD and homo-tri-RBD. **a.**Native structure of S protein trimer. Cyan, red and green colors represent three RBDs which were assembled into a trimeric arrangement. **b.**Structural modelling of the homo-tri-RBD protein by Modeller software using the native structure of S protein as the template. **c.**Structural modelling of the mutI tri-RBD protein by Modeller software using the native structure of S protein as the template.

**Extended Data Fig. 2.**
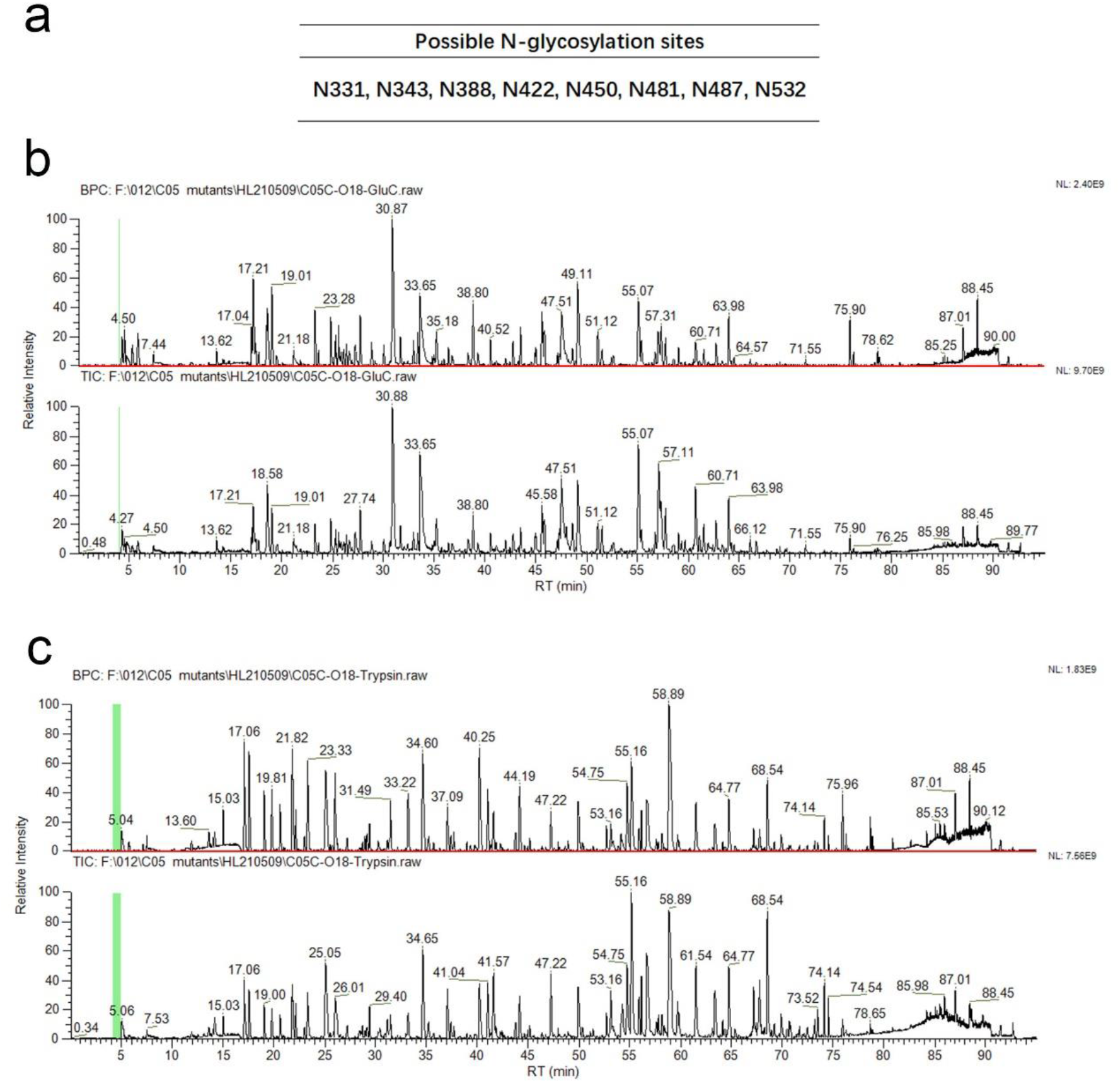
Base peak chromatogram (BPC) and total ion Chromatogram (TIC) to identify the N-glycosylation sites by UPLC-MS. **a.**The identified possible N-glycosylation sites for one RBD unit in mutI tri-RBD. **b.**BPC (upper subfigure) and TIC (lower subfigure) for mutI tri-RBD protein digested by GluC to identify N-glycosylated sites. **c.**BPC (upper subfigure) and TIC (lower subfigure) for mutI tri-RBD protein digested by Trypsin to identify N-glycosylated sites.

**Extended Data Fig. 3.**
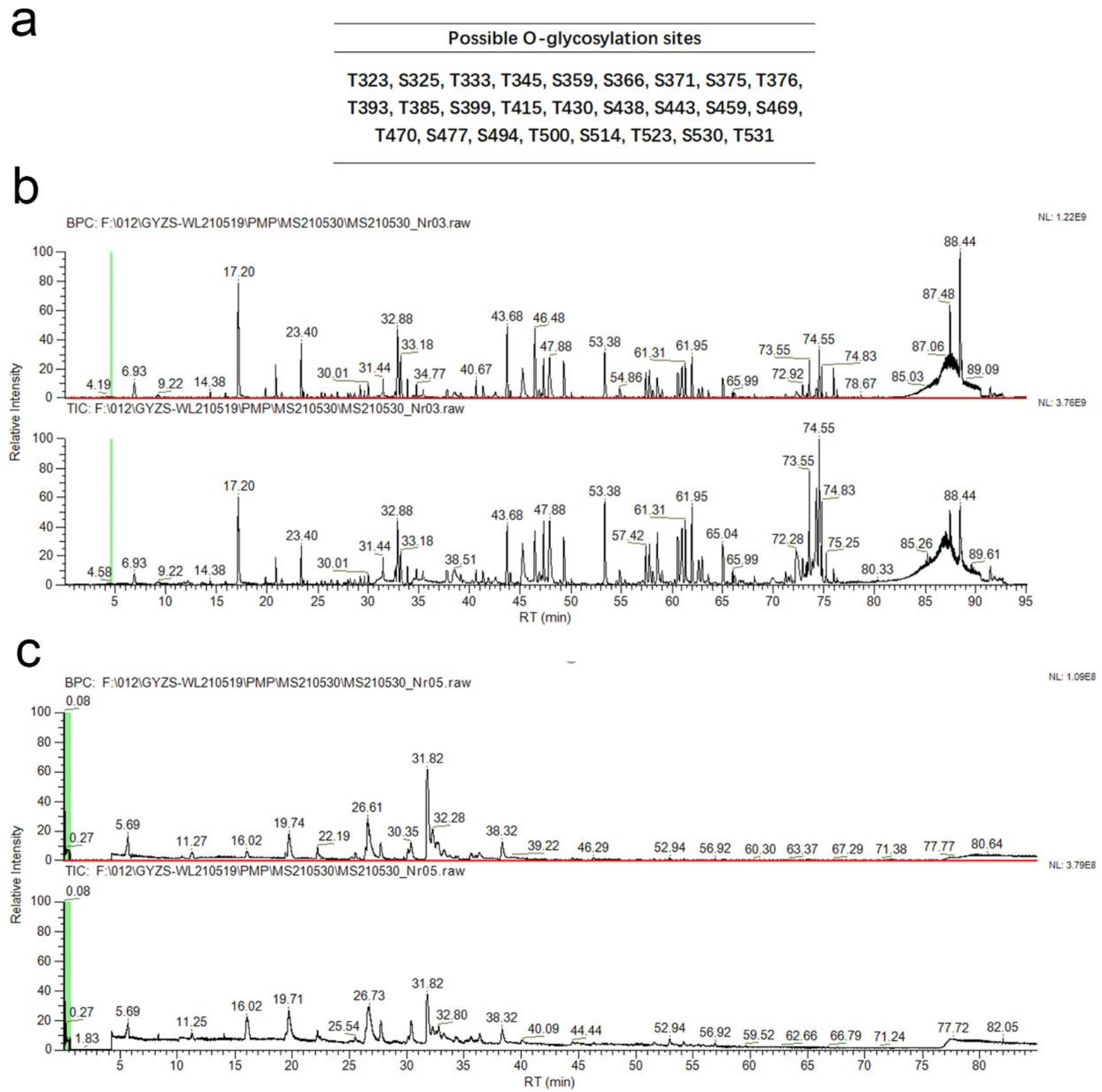
Base peak chromatogram (BPC) and total ion Chromatogram (TIC) to identify the O-glycopeptides by UPLC-MS. **a.**The identified possible O-glycosylation sites for one RBD unit in mutI tri-RBD. **b.**BPC (upper subfigure) and TIC (lower subfigure) of digestion product to identify larger O-glycosylated peptides. **c.**BPC (upper subfigure) and TIC (lower subfigure) of the purified smaller peptides to identify O-glycopeptides.

**Extended Data Fig. 4.**
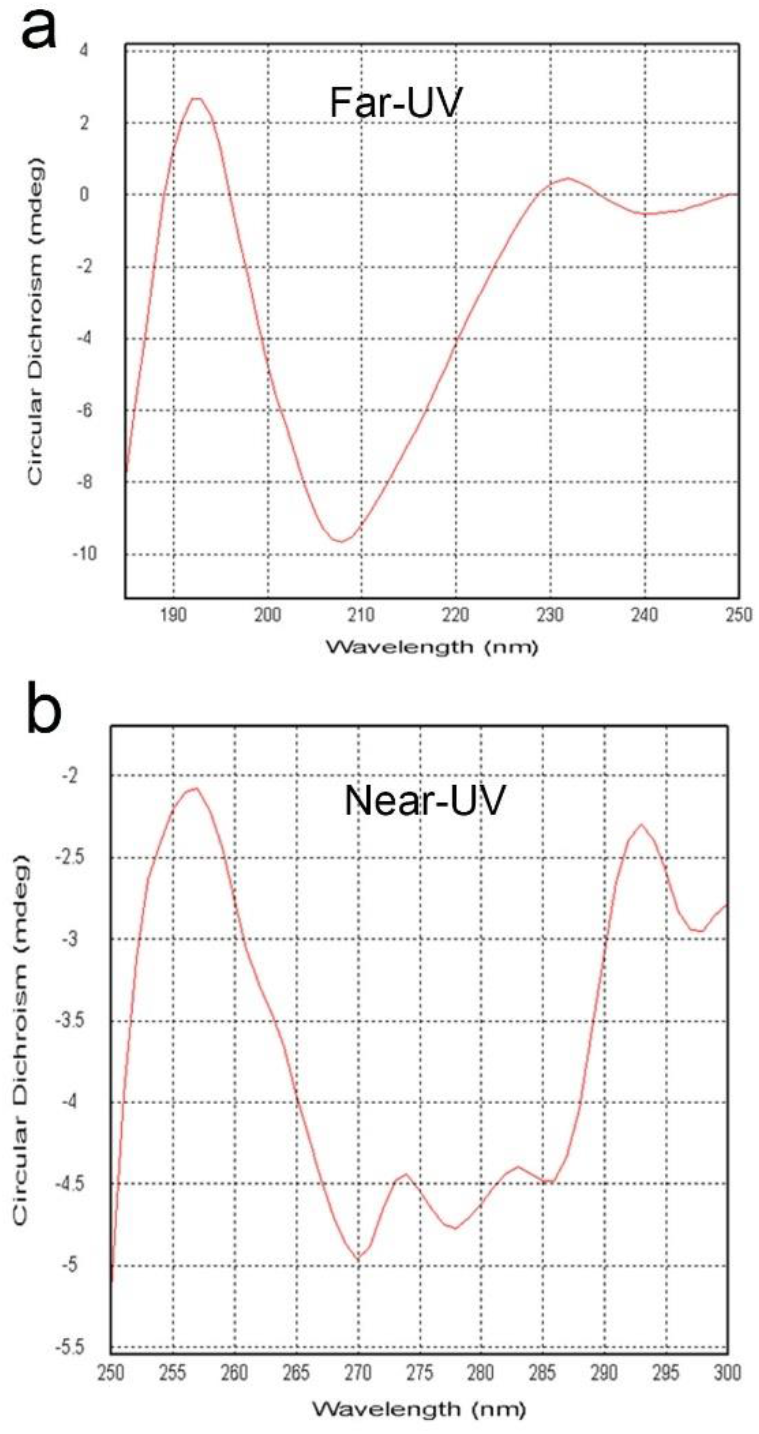
Far-ultraviolet (UV) and near-UV spectra acquired by Circular Dichroism (CD) to estimate the content of different secondary structures in the recombinant mutI tri-RBD. **a.**Far-UV spectra acquired by CD. **b.**Near-UV spectra obtained by CD.

**Extended Data Fig. 5.**
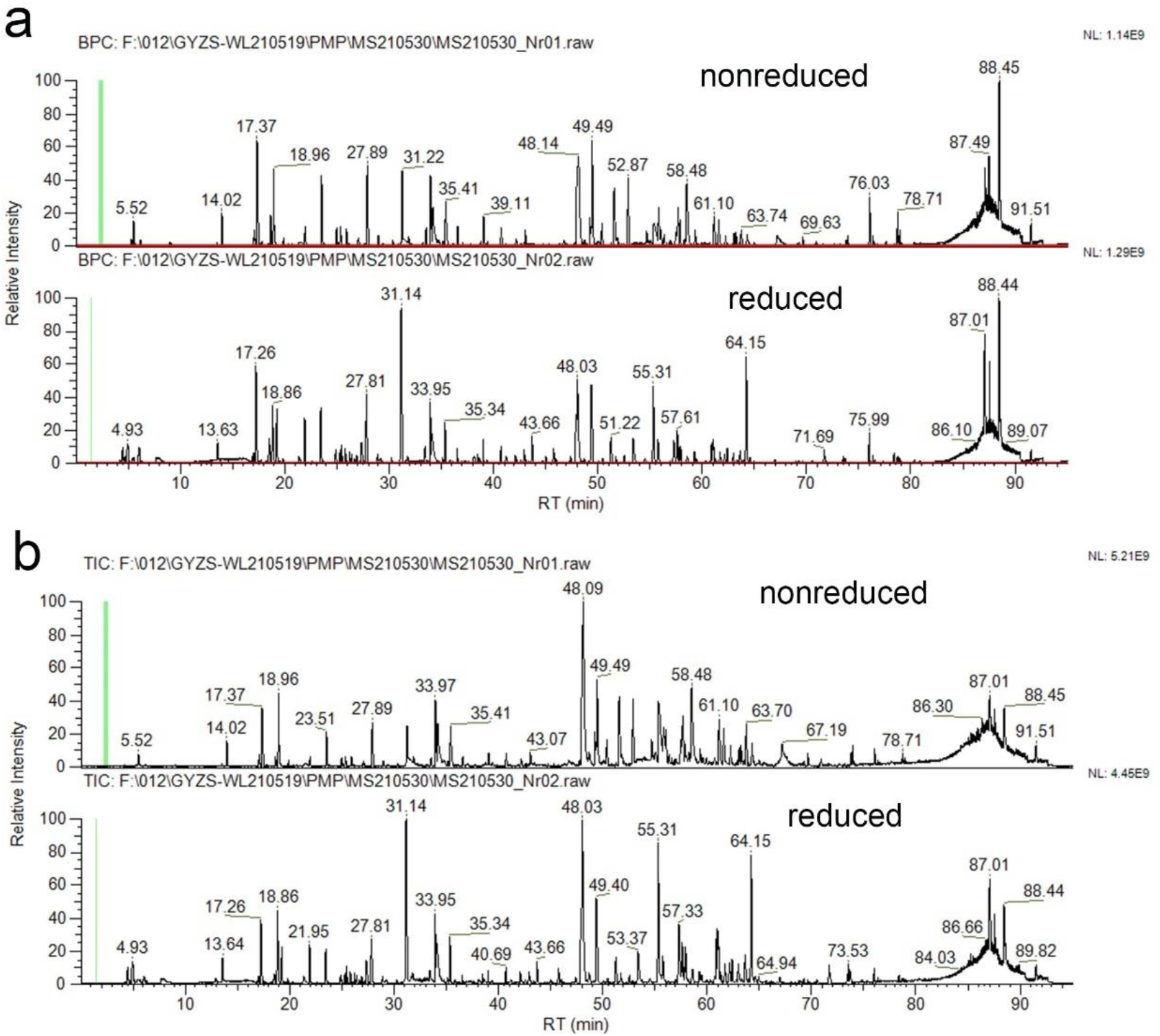
Base Peak Chromatogram (BPC) and Total Ion Chromatogram (TIC) to identify the disulfide-bonds in mutI tri-RBD by UPLC-MS. **a.**BPC for the nonreduced and reduced protein sample. **b.**TIC for the nonreduced and reduced protein sample.

**Extended Data Fig. 6.**
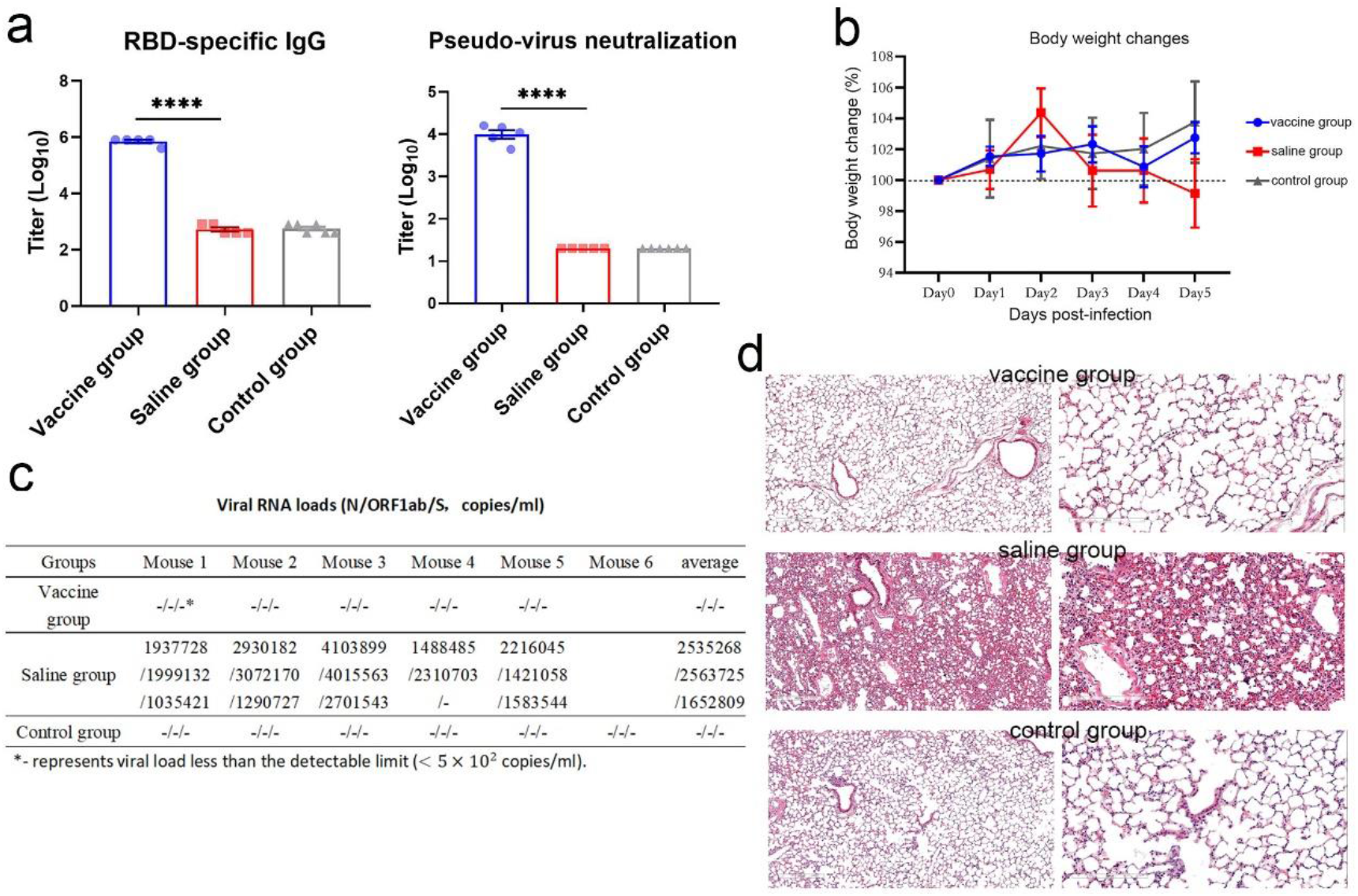
Protection of homo-tri-RBD vaccine candidate against live SARS-CoV-2 prototype virus challenge in transgenic mice. The mouse challenge experiments were performed in the ABSL3 facility. All the animal experimental procedures were approved by the Institutional Animal Care and Use Committee of the National Vaccine and Serum Institute of China, and Institutional Animal Care and Use Committee of National Institute for Viral Disease Control and Prevention, Chinese Center for Disease Control and Prevention, Beijing, China. A total of 16 female ACE2-TG/IOZ transgenic mice aged 6-10 weeks (purchased from Beijing Huafukang Bioscience Co., Ltd, China) were used for the live SARS-CoV-2 challenge experiment, which were randomly divided into 3 groups: a vaccine group (5 mice), a saline group (5 mice) and a control group (6 mice). The mice in vaccine group were immunized with two intramuscular injections of 2 μg/dose candidate vaccine on Day 0 and Day 21. In saline group and control group, the mice were intramuscularly injected with physiological saline on Day 0 and Day 21. On day 7 after the second vaccination, the blood from the caudal vein of the immunized mice was collected, and serum was separated to measure the titers of specific IgG and neutralizing antibodies using ELISA and pseudo-virus neutralization assay, respectively. **a.**The titers of specific IgG and pseudo-virus neutralizing antibodies in the sera of the immunized mice compared with those of the mice in the saline and control groups. Data are presented as mean ± SEM. P values were determined by Student’s t-test. On day 14 after the second immunization, the mice in vaccine group (5 mice) and saline group (5 mice) were challenged with 1.5 × 10^5^ *TCID*_50_ of live SARS-CoV-2 prototype virus via the intranasal route. The mice in the control group were not subjected to live virus challenge, serving as the blank control. During the challenge experiment, the body weight of the mice was measured every day. **b.**Comparison of the body weight changes of the mice in the three groups during the challenge experiment. Five days after virus challenge, all mice were euthanized and the lung tissues were collected. The viral RNA was extracted from lung tissues by using a nucleic acid extraction kit (QIAGEN, No. 52906), and the viral load was evaluated by using a 2019-nCoV (N, ORF1ab and S genes) nucleic acid detection kit (PCR-fluorescence probing) purchased from Mabsky (Shenzhen) with product No. 349. **c.**Comparison of the viral loads in the lung tissues of the mice in the three groups. N, ORF1ab and S viral genes were detected. In addition, the histopathological sections of the lung tissues of the mice were stained with hematoxylin and eosin (HE), and the pathological examination was performed. **d.**Histopathological examinations of the lung tissues of the mice in the three groups. Left subfigure in each group displays the lung tissue amplified four times, and right subfigure enlarged ten times. Live virus challenge experiment showed that on day 7 after the whole vaccination, high titers of specific IgG and neutralizing antibodies were detected in all mice immunized with homo-tri-RBD vaccine candidate, whereas in the saline and control groups, no antibody response was detected (see panel **a**). During the live virus challenge experiment, the body weight of the mice in the saline group also obviously reduced, suggesting possible infections of SARS-CoV-2. While for the mice in vaccine group, the body weights fluctuated within the normal range, which is similar to the mice in control group (see panel **b**). This result indicated that homo-tri-RBD candidate vaccine might provide protections against the infection of live SARS-CoV-2. The viral load detections in the lung tissues showed that all the mice in saline group exhibited very high viral loads after live virus challenge, and the average viral load of the five mice reached 2535268 copies/ml, 2563725 copies/ml and 1652809 copies/ml for N, ORF1ab, and S genes, respectively (see panel **c**). In contrast, the viral loads in lung tissues for all the mice in the vaccine group were under the detectable limit (< 5 × 10^2^ *copies/ml*), which implies that our candidate vaccine could provide complete protections to the transgenic mice against the live SARS-CoV-2. The histopathological examination of the lung tissues showed that severe interstitial pneumonia, a typical feature of COVID-19, occurred in the mice of the saline group (see panel **d**). Substantial histopathological changes in the lung tissue were observed, including alveolar septa thickening, heavy inflammatory cell infiltration and focal hemorrhage. Large amounts of exudates appeared in lung cavities, as well as the formation of hyaline membrane and bullae were also observed. In addition, tracheal bronchus and blood vessels also exhibited a severe infiltration and exudates combined with desquamation of epithelial cells. In contrast, no obvious histopathological change was observed in the lung tissues of the mice received vaccination, indicating our candidate vaccine could effectively block the infections of SARS-CoV-2 in the transgenic mice.

